# Improved spatial memory in a modular network mimicking the prefrontal-thalamo-hippocampal triangular circuit

**DOI:** 10.64898/2026.02.27.708561

**Authors:** Munenori Takaku, Tomoki Fukai

## Abstract

The hippocampus (HPC), prefrontal cortex (PFC), and thalamic nuclei, such as reuniens (Re), form a reciprocally connected circuit that plays a critical role in processing hippocampus-dependent memory. Accumulating evidence suggests that this triangular modular circuit is crucial for performing cognitive tasks that require context-dependent memory, which belong to a class of behavioral tasks difficult for animals to learn. Experiments are gradually revealing what behavioral information these brain regions represent, but how the triangular circuit gives rise to the observed divisions of labor remains unknown. It is also unclear whether the triangular modular circuit brings any advantage in solving such tasks. Here, we addressed these questions by constructing a prefrontal-thalamo-hippocampal circuit model comprising interconnected long-short-term memory (LSTM) units and training it on contextual memory-dependent spatial navigation tasks. Our model revealed the critical roles of the distinct brain modules. The HPC module encoded spatial information, whereas the PFC module represented the spatiotemporal task structure in a context-dependent manner. The Re module integrated task-relevant information to facilitate learning in the PFC and HPC modules, dynamically harmonizing these modules. The thalamic coordination of the other modules enhanced the system’s robustness in learning to navigate complex environments. This division of labor between the HPC, PFC, and Re modules was not specified a priori but emerged through learning, showing an interesting coincidence with the task-related activities of the prefrontal-thalamo-hippocampal circuit. Our results demonstrate that the multi-modular network structure is crucial for robust processing of context-dependent memory.

## Introduction

Understanding how the brain stores and recalls memories about an episode is a fundamental question in neuroscience. The prefrontal cortex (PFC) and hippocampus (HPC) are the locus of storing task-relevant information when subjects solve memory-dependent tasks subject to context-dependent rules [1]. During cognitive tasks, these regions exhibit distinct yet complementary encoding patterns [2– 5]: hippocampal neurons primarily encode spatial and positional information [6], while prefrontal cortical neurons represent contextual rules of decision behavior and the resultant value changes [7, 8]. However, the coordination of multiple brain regions to harmonize different cognitive computations is poorly understood.

The thalamus has been shown to influence both hippocampal and prefrontal functions. This region not only sends sensory information to the cortex and gates sensory input [9], but also participates in conducting various cognitive tasks [10]. Thalamic lesions in rodents cause spatial memory impairment and loss of temporal order, [11–13], indicating thalamic involvement in cognitive processing. The ventral midline thalamic nuclei, including the nucleus reuniens (Re) and rhomboid, occupy a position that may influence the HPC-PFC network as a whole. These nuclei represent a major source of thalamic input to the hippocampus and maintain connections with the medial PFC [14, 15]. Research has been increasing on the structure and function of the PFC-thalamo-HPC circuit [16–19]. Notably, inactivation of the ventral midline thalamus selectively impairs performance on behavioral tasks that require PFC-HPC interactions [20, 21]. Impairment of the ventral midline thalamus, particularly Re, is also implicated for disrupted PFC-HPC coordination, including changes in their phase synchrony and coherence across multiple frequency bands [22–24]. These observations suggest that the thalamic activity is highly engaged in the cooperation between PFC and HPC during memory processing. The triangular architecture of the PFC-thalamo-HPC circuit may therefore be crucial for performing contextual memory-dependent cognitive tasks.

Despite accumulating evidence about the behavioral information these brain regions represent, critical questions remain to be explored. How do the three regions cooperate in processing various kinds of information about episodes in different behavioral environments? How do they enable robust processing of context-dependent memory? What advantages might this triangular modular architecture offer over simpler circuits? Here, we addressed these questions by constructing a PFC-thalamo-HPC circuit model comprising interconnected Long Short-Term Memory (LSTM) units [25, 26]. Instead of modeling complex cortical and thalamic circuits, we employed LSTMs for their capacity to implement gating functions analogous to those played by cortical inhibitory cells and thalamic modulation. A circuit model of LSTMs was previously used to account for reward-driven sequence learning in the PFC [27]. We trained our model on context-dependent spatial navigation tasks that rely on working memory in a context-dependent manner, a class of behavioral tasks difficult for animals to learn [28].

Our model revealed that distinct functional roles emerge in the three regions (modules) through learning: the HPC module encodes spatial information, the PFC module represents context-dependent task rules, and the Re module integrates the task-relevant information while coordinating the other modules. This division of labor was not predetermined but emerged through learning and is consistent with biological observations. Furthermore, we compared task performance between different three-module architectures, which revealed that the biologically inspired architecture has some computational advantages over the other architectures, particularly in learning speed and robustness in learning complex tasks. Our results demonstrate how the biology-inspired structure and functional specializations of the multi-module PFC-Re-HPC circuit benefit the robust processing of context-dependent memory.

## Results

### Three-module model and task settings

For convenience in terminology, hereafter, we call the thalamic circuit in our model “Re module”. We developed a computational model consisting of three specialized modules representing the HPC, PFC, and Re (Fig 1A). The architecture of this model was inspired by the anatomical connectivity among these brain regions and enabled us to study their functional interactions during memory-dependent navigation tasks. The HPC and PFC modules comprise 20 recurrently connected Long Short-Term Memory (LSTM) units each [25, 26]. Each LSTM unit contains input, output, and forget gates along with a memory cell that uses sigmoidal activation functions. The Re module was implemented as an Elman network of 20 recurrently connected units without gating units. The module sizes were chosen to balance between the model’s complexity and learning capability (See the Methods).

**Fig 1.**
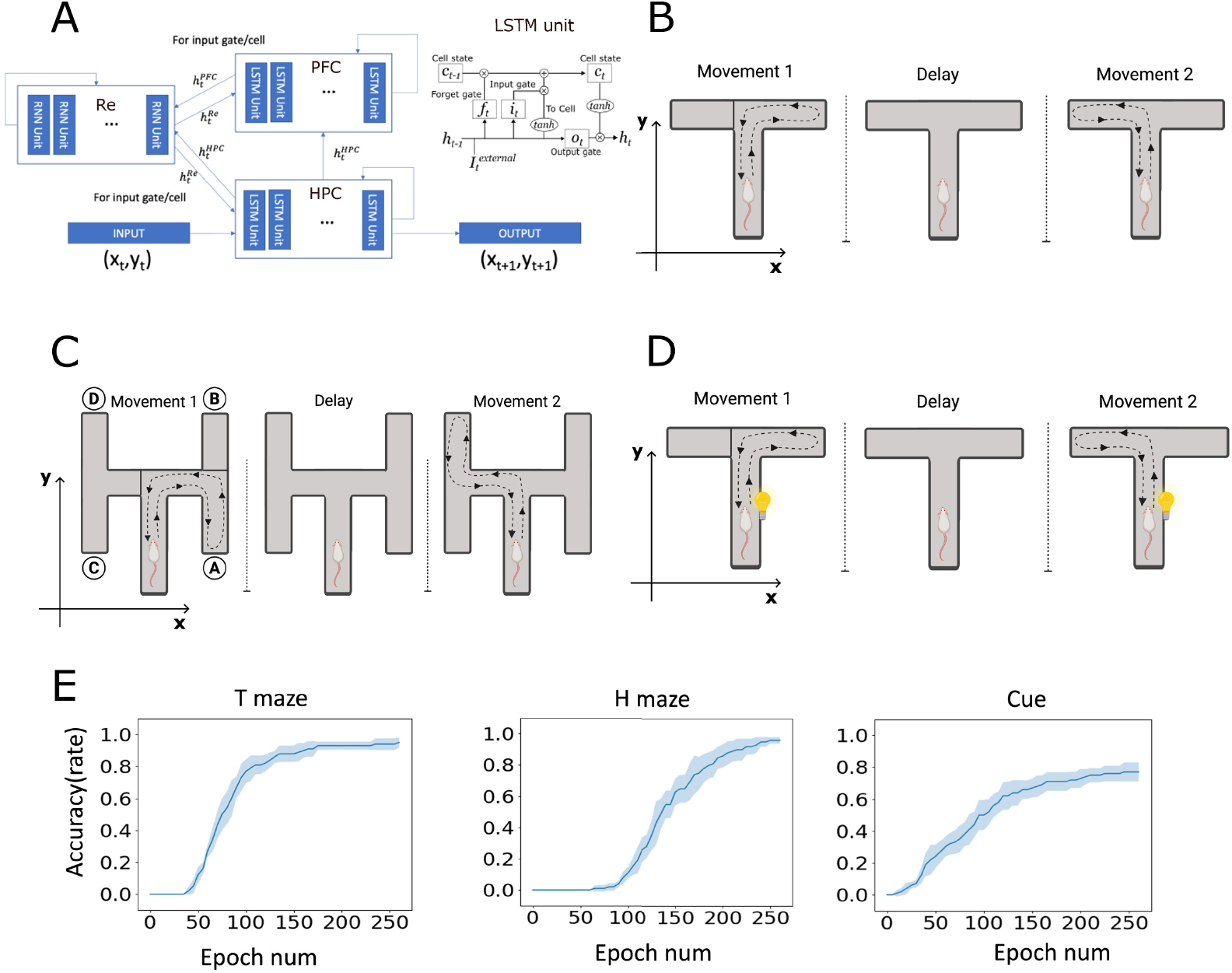
Architecture of the three-module model. B-D panels are created in https://BioRender.com. **A**. The model consists of HPC and PFC modules implemented as LSTM networks (inset) and the Re module as an Elman network without gating functions. The model has bidirectional connections between the Re and HPC/PFC modules and a directed projection from HPC to PFC. The HPC module receives an external input and outputs the agent’s current location. **B**. A delayed non-match to sample (DNMS) task on a T-maze. The agent should first visit one arm of the T-maze, return to the home position (Movement 1), stay there during a fixed delay period (Delay), and then visit the opposite arm (Movement 2). **C**. A DNMS task on an H-maze. Unlike the T-maze task with two round trips, the H-maze task involves a fixed sequence of four round trips between the home position and positions A, B, C, and D. After returning to the home position, the agent is required to stay there during a fixed delay period. **D**. A DNMS task with cue signals. A brief binary cue was added at the end of a delay period in the T-maze task to indicate the time to leave the home position. **E**. Learning curves of the three-module model are shown for the three tasks. Shaded areas show SD.

Information flow within the network follows a specific pattern, where the HPC module receives sensory input and generates predictions of subsequent states, projecting to both the PFC and Re modules. The PFC module receives input from the HPC module and communicates with the Re module, which then integrates inputs from the HPC and PFC modules and projects back to both of them. Importantly, we restricted the Re-to-HPC/PFC feedback to their input gates and memory cells, assuming that thalamic signals modulate inputs to cortical neurons but do not gate their outputs.

We evaluated our model on a delayed non-match to sample (DNMS) task [1], which is a spatial decision-making paradigm widely used in rodent studies. This task requires processing of spatial information with temporal context and working memory ability. We tested the model for three task settings: a T-maze case, an H-maze case, and a cue-applied case (Fig 1B, C, D). In the T-maze condition, the model navigates a T-shaped environment where the agent must first visit one arm, return to the starting position, stay there during a delay period, and then navigate to the opposite arm to receive a reward. Each trial consisted of 100 discrete time steps. In the H-maze condition, we designed a more complex environment featuring four potential arms, requiring the model to learn specific sequential arm-visit patterns consisting of 160 discrete time steps per trial. In the cue-applied condition, we modified the T-maze paradigm by providing external cue signals at the trial initiation and the termination of the delay period, with variable trial durations ranging from 80 to 120 time steps.

Context is defined as latent environmental factors that influence animal’s behavior other than observable sensory information. In the present DNMS task, different contexts arise within the same spatial environment from different decision rules and the temporal task structure. To make successive choices correctly, the agent has to maintain the contextual information internally during the task period. We investigate how the dynamics of the prefrontal-thalamo-hippocampal network model establish such internal states.

For the T-maze task, successful performance was defined as alternating between two maze arms (Fig 1B) at least three times in a correct sequence. For the H-maze task, the model needed to reach specific arm coordinates three times in the prescribed order of A→D, B→C, C→A, and D→B (Fig 1C). A model is considered “Good” when it reproduces the target trajectory accurately enough. A model was classified as “Failed” if it could not leave the initially visited arm or exhibited repetitive non-goal-directed movements. The accuracy metric refers to the proportion of Good models among 100 independently trained models. After training, the three-module model exhibited proficient performance across all task variations (Fig 1E), suggesting that the biologically inspired network architecture can support memory-dependent spatial navigation tasks with varying complexities and difficulties. Thus, the model is flexibly generalizable to a diverse range of spatial decision-making tasks.

### Distinct representations of task-related information in the three modules

We conducted principal component analysis (PCA) to investigate the low-dimensional dynamical features of each processing module. PCA revealed interesting differences between the three modules. Unlike the other modules, the activity of the HPC module was well explained by the first principal component (PC1), accounting for approximately 60% of the variance (Fig 2A). In contrast, the PFC and Re modules required multiple components (up to PC4) to explain 60%-70% of their activities, indicating more complex, multidimensional representations. A critical difference is visible during delay periods within the PC1: the variation gradient of the HPC activity vanished rapidly, while the gradients of the PFC and Re modules remained finite (Fig 2B). The vanishing PC1 gradient of the HPC module implies that it represents such information that remains unchanged during delay periods, namely, the spatial location of the agent. In contrast, the PFC and Re modules represent such information that varies dynamically during the entire task period and hence extends beyond purely spatial information.

**Fig 2.**
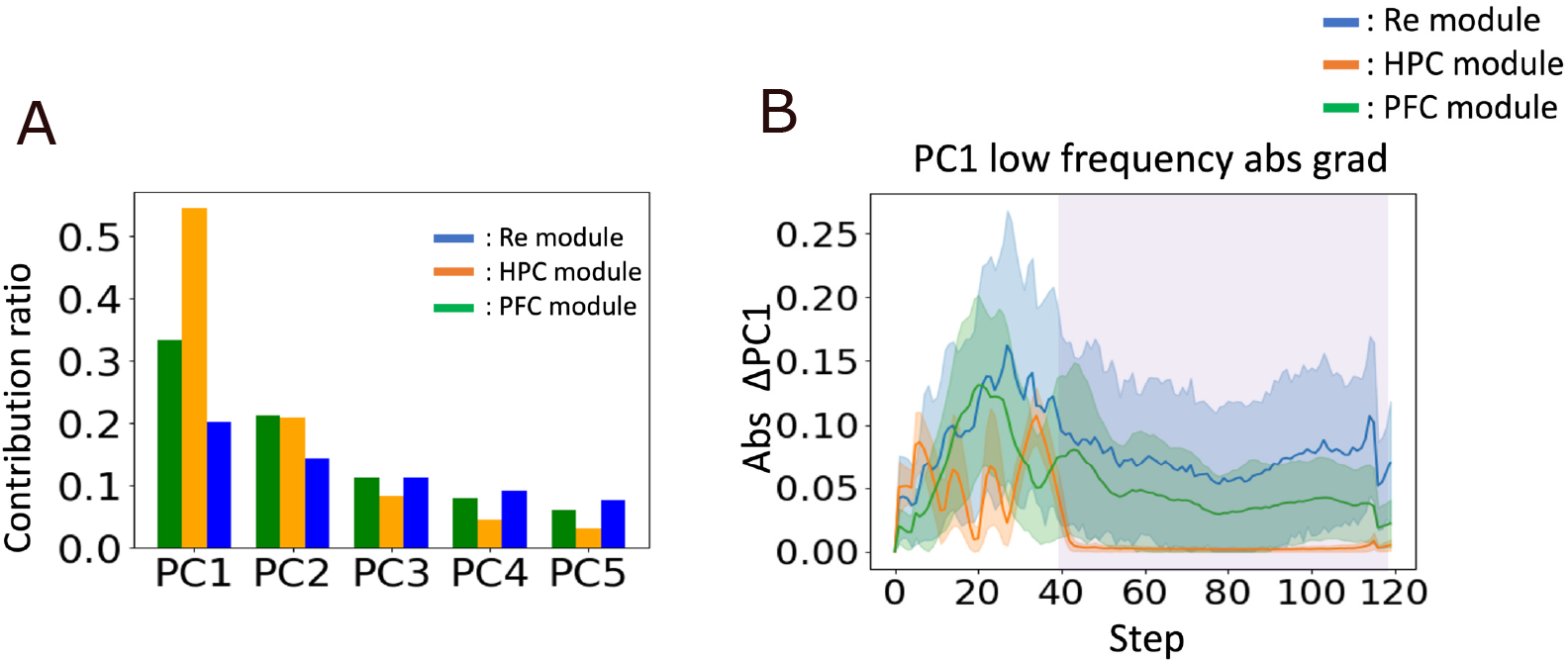
Principal component analysis of module activities. **A**. The contribution ratios of the first five principal components (PC1-PC5) are plotted for each module. The ratios were averaged across 100 trained models. **B**. The temporal gradients of low-pass filtered PC1 values during task execution are shown for the three modules. The gradients represent the absolute values of the rate of change smoothed with a 5-step moving average filter. Error bars represent SD. The results were averaged across all successful trials of 100 trained models.

We further conducted PCA for the network model trained on the H maze. We first compared the dynamical behavior of the leading principal components between the HPC and PFC modules (Fig. 3A, B). In the HPC module, the leading components, PC1 and PC2, represented changes in the agent’s location during the spatial navigation task, with the shape of the learned neural trajectories directly reflecting that of the maze. In particular, the agent’s locations along the four different arms were represented by combinations of PC1 and PC2 (Fig. 3C). Furthermore, these components remained constant during the delay period (Fig. 3C), which is consistent with spatial information coding in the HPC module. The next leading-order components, PC3 and PC4, exhibited fluctuation around 0, presumably reflecting small trial-by-trial variations in the agent’s trajectories.

**Fig 3.**
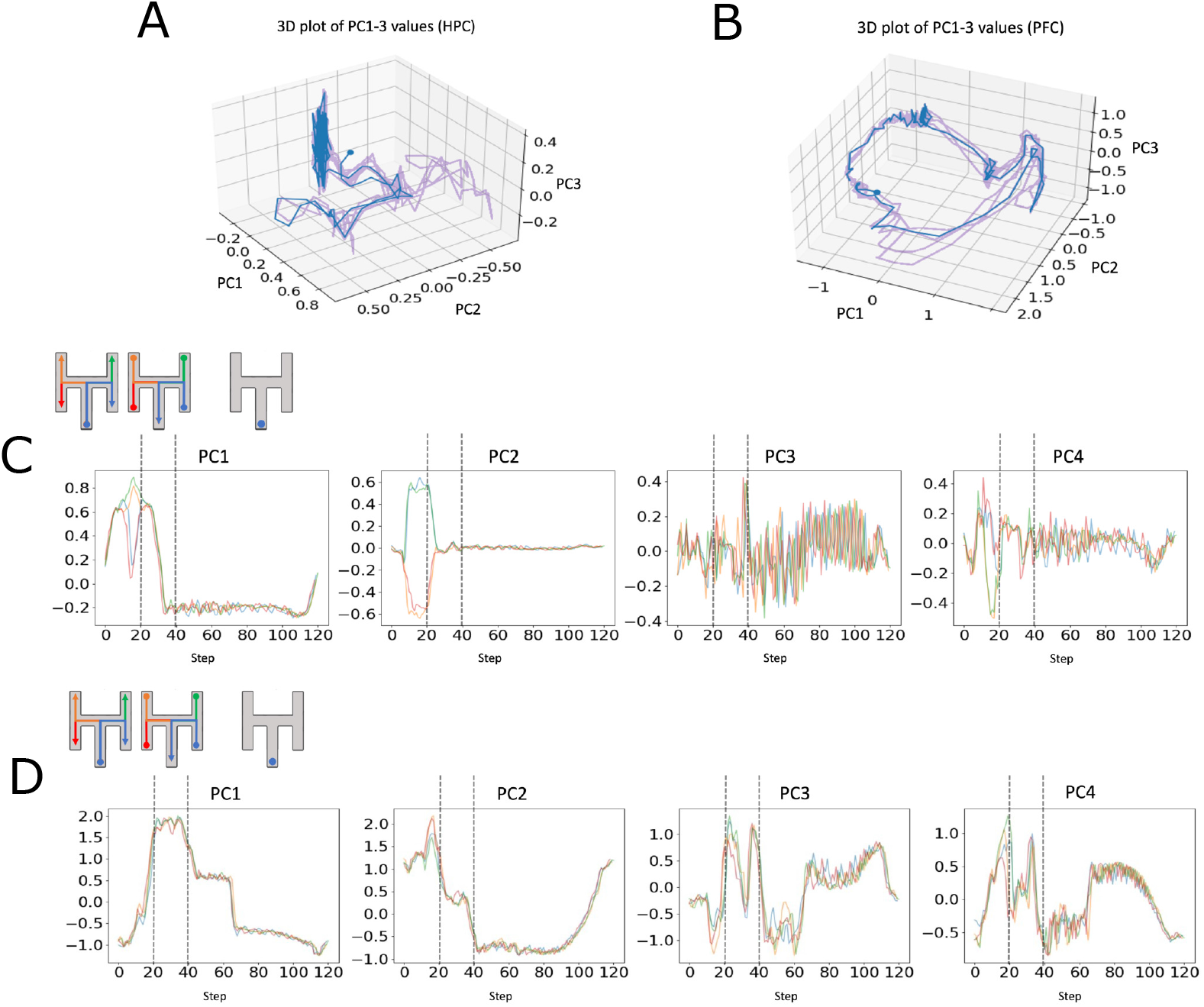
Representations learned by the HPC and PFC modules. The results are shown for the H-maze task. **A**. Three-dimensional PCA plot of HPC module activity. The blue-colored portion of the entire trajectory (purple) corresponds to right-bottom arm visits. **B**. Three-dimensional PCA plot of PFC module activity shown with the same color scheme as in A. **C**. The temporal evolution of the first four principal components in the HPC module. Trajectories visiting four arms (top) and the corresponding principal components (bottom) are marked with the same colors. Vertical dotted lines divide the task into three behavioral phases: arm visit, return to the home position, and delay period. **D**. The temporal evolution of the first four principal components is shown for the PFC module in the same format as in C.

Contrastingly, the leading principal components of the PFC module lacked clear structures that represented spatial information (e.g., the locations of the four arms) but exhibited a multifaceted structure (Fig. 3B). For example, the leading components, PC1 and PC2, discriminated different behavioral phases of the task, such as the epochs of movement and waiting still. During the delay period, these components behaved as if they had measured the timing of the next movement, although they did not discriminate between visits to different arms of the maze (Fig. 3D). Interestingly, visits to different arms are more clearly distinguishable in PC3 and PC4 than in PC1 and PC2 during both movements and the delay period. Therefore, the PFC module represents spatial information through sub-leading principal components. These results suggest that the PFC module likely processes the global features of the task, such as behavioral phases, temporal organization, and decision rules, that are represented over many principal components.

From these result, the HPC module’s gradient (corresponding to spatial information) reached zero during delays, while the Re module’s gradient did not, indicating that the Re module encodes information beyond purely spatial representations (Fig 2B). The observed differentiation in representational content suggests that the model acquires this specialization of functional roles through training, potentially reflecting a systems-level specialization that enhances task optimization and generalization.

### The roles of Re Module

The principal components of the Re module exhibited characteristic fluctuations, which appear to encode and integrate various types of task-related information. First, the leading component (PC1) displayed slight differences between visits to the left arm and right arm (Fig 4A), indicating that this module also represents spatial information. Second, as in the PFC module, the Re module showed gradually increasing or decreasing activity during delay periods, which likely coordinates the onset timing of the agent’s spatial movements (Fig 4A). Third, the fluctuations of PC1 displayed a consistent difference between left-arm and right-arm visits during delay periods (Fig 4B). Since the agent’s spatial location did not change significantly during the delay periods, the difference in the principal component likely encoded information about a rule to decide between the left arm and right arm in the next trip.

**Fig 4.**
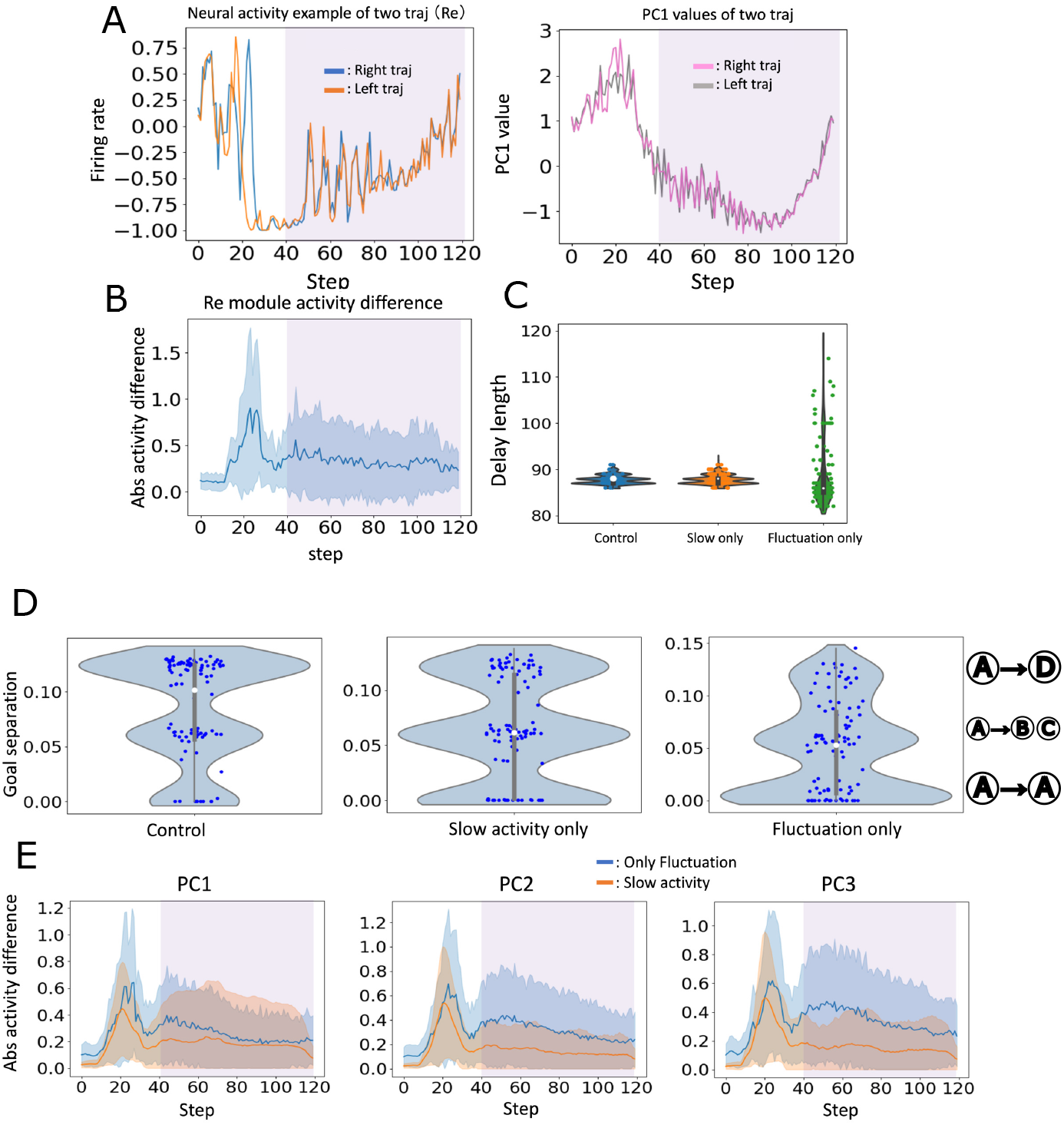
Representations learned by the Re module. All results are shown for the H-maze task. Shaded purple areas show the delay period. **A**. Left: Representative neural activity from the Re module. The shaded area shows the delay period. “Right traj” and “Left traj” indicate trajectories visiting positions A and D, respectively. Right: The PC1 component of the Re module activity. Two portions highlighted (red circles) demonstrate slight differences between left and right trajectories during movement and delay periods. **B**. The average difference (thick line) of PC1 values between left and right trajectories is shown. The shaded area indicates SD. **C**. The delay periods learned are shown for the control and two truncated models. The target delay length was set to 86–90 steps, allowing a small range for technical reasons. **D**. Distances in the x-coordinate between the initial and final positions of individual trips (indicated at right) are displayed to visualize errors in navigation. For instance, plots with near-zero distances correspond to error trips in which the agent visited the starting arm. **E**. The time evolution of the leading principal components during the entire task period is shown for the two models.

The Re module was crucial for integrating spatial, temporal, and contextual information about the spatial decision-making task. To examine how temporal information is represented in the Re module, we analyzed the delay length of the learned trajectories under two conditions. To this end, we applied a moving-average filter to these fluctuating trajectories to extract their low-frequency components (slow changes). In the “Fluctuation-only” condition, we replaced the output of the Re module with the high-frequency components, eliminating the slow component of the Re module activity from the model. In contrast, in the “Slow-only” condition, we replaced the output of the Re module with the slow component, eliminating the high-frequency components of the Re module activity. The delay length remained unchanged in both the normal and Slow-only conditions, whereas it varied significantly in the Fluctuation-only condition (Fig. 4C). These results demonstrate that temporal information about the delay period was primarily encoded in the slow component of the fluctuating Re activity.

We analyzed error trials to find that the fluctuating components are essential to enable the agent to switch between learned behaviors. Figure 4D) shows the distances from the initial point (arm A) to the end points the agent traveled in error movements (hence the agent failed to return to the home position) in the three conditions. Almost vanishing distances for A→ A indicate that the agent failed in leaving the initial location. When only the slow component was supplied to the model, the agent failed to leave the initial arm more often than in the control condition, as indicated by increases in the travels A→A and A→ B. When only high-frequency fluctuating components were supplied, the agent’s behavior became fragile, and it failed to replicate the learned trajectory. These results suggest the crucial role of the fluctuating components in generating appropriate transitions between the learned state representations.

We further investigated the roles of the fluctuating components by performing PCA of the Slow-only and Fluctuation-only models (Fig 4E). The PC1 components of the two models showed similar behavior during spatial exploration, suggesting that spatial information is represented in both frequency domains. During the delay period, the PC1 of the two models behaved similarly, but the Fluctuation-only model dominated the Slow-only model in PC2 and PC3. These results suggest that the contextual information (about the decision rules) of the task is preferentially encoded in the high-frequency components. Thus, the Re module employs a frequency-multiplexing code: while spatial information for behavioral stability is distributed across frequencies, temporal information and contextual information governing behavioral switching are represented in the slow component and high-frequency fluctuation of RE activity, respectively.

Another important role of the fluctuating dynamics of the Re module was to facilitate synchronization between the HPC and PFC modules through its reciprocal connections with these modules (Fig 5A). To measure this synchronization, we calculated coherence between module activities using spectral analysis. Specifically, we applied short-time Fourier transforms with a window width of 60 steps to generate spectrograms for each module and then calculated cross-spectra for the Re-HPC and Re-PFC module pairs. Our analysis revealed that synchronization emerges progressively through training in both Re-HPC and Re-PFC couplings (Fig 5B, C). Well-trained, successful models exhibited significantly higher coherence values compared to poorly-trained, failed models (Fig 5D: *p*<0.05 for Re-PFC; *p*<0.001 for Re-HPC, one-way ANOVA). These results demonstrate the Re module’s influence on network-wide synchronization, intriguingly aligning with similar experimental findings in the nucleus reuniens [18]. Thus, the Re module synchronizes the HPC module and PFC module, and represents contextual, spatial, and temporal information in an integrated manner. If the three modules cannot cooperate synchronously enough, the entire model fails to learn the memory-guided spatial decision-making task.

**Fig 5.**
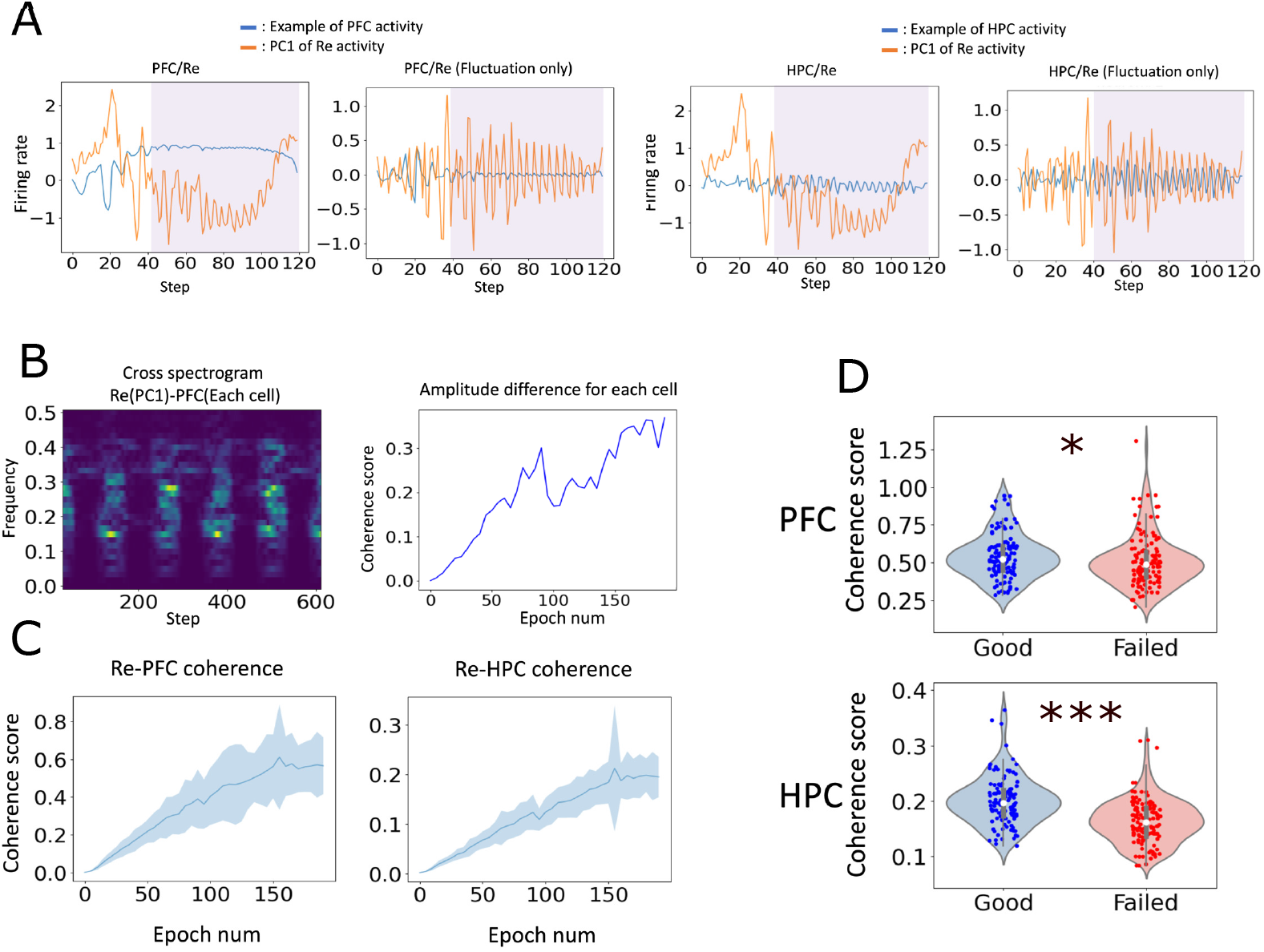
Synchronization between the Re and PFC/HPC modules. Shaded purple areas show the delay period. **A**. Examples of oscillating PFC/HPC activities are shown together with the PC1 of Re activity in the control and fluctuation-only conditions. **B**. Left: The cross-spectrogram between the Re and PFC modules was calculated to show behavioral phase- and frequency-specific Re-PFC synchronization patterns. Right: The evolution of pair-wise synchronization strength between these modules is shown during learning. **C**. The evolution of the average coherence scores (solid lines) between the Re and PFC(left)/HPC(right) modules is shown during learning. Shaded areas show SD. **D**. The degrees of synchronization were compared between successful (‘Good’) and unsuccessful (‘Failed’) learning models. Successful models exhibited significantly higher scores between the Re-PFC (p<0.05) and Re-HPC (p<0.001) modules.

Finally, we examined the contribution of the Re module to learning by removing it from the model (S1 FigA). Both HPC and PFC modules consisted of 30 LSTM units, such that the Re-truncated model has the same total number of LSTM units as the original model, which has 20×3 = 60 LSTM units. Additionally, we introduced PFC-to-HPC feedback connections to compensate for the loss of the synaptic pathway from the PFC to the HPC through the Re module. We trained the Re-truncated model on the T-maze and H-maze tasks, and a cue-triggered T-maze task. In all three cases, the original 3-module model outperformed the Re-truncated model. In particular, the 3-module model performed much better than the Re-truncated model in the H-maze and cue-guided tasks (S1 FigB). These results indicate the crucial contribution of the Re module to learning spatial decision-making tasks, particularly in complex conditions, presumably through the coordination of the HPC and PFC modules by the Re module.

### The origins of individual variances in learning performance

Our model exhibited large individual variances in learning the present task. To elucidate what distinguishes well-trained models from poorly trained ones, we compared the eigenvalue spectra of weight matrices between the two categories of models. Because both categories of models learned spatial representations equally well, we investigated the structural differences between their PFC modules. First, we analyzed the eigenvalue spectra of recurrent connections between early and late stages of learning (Fig 6A). During learning, the average lengths of complex-valued eigenvalues rapidly grew. Notably, the average eigenvalue length was significantly larger in well-trained, successful models than in poorly trained, failed models (*p*<0.01, ANOVA).

**Fig 6.**
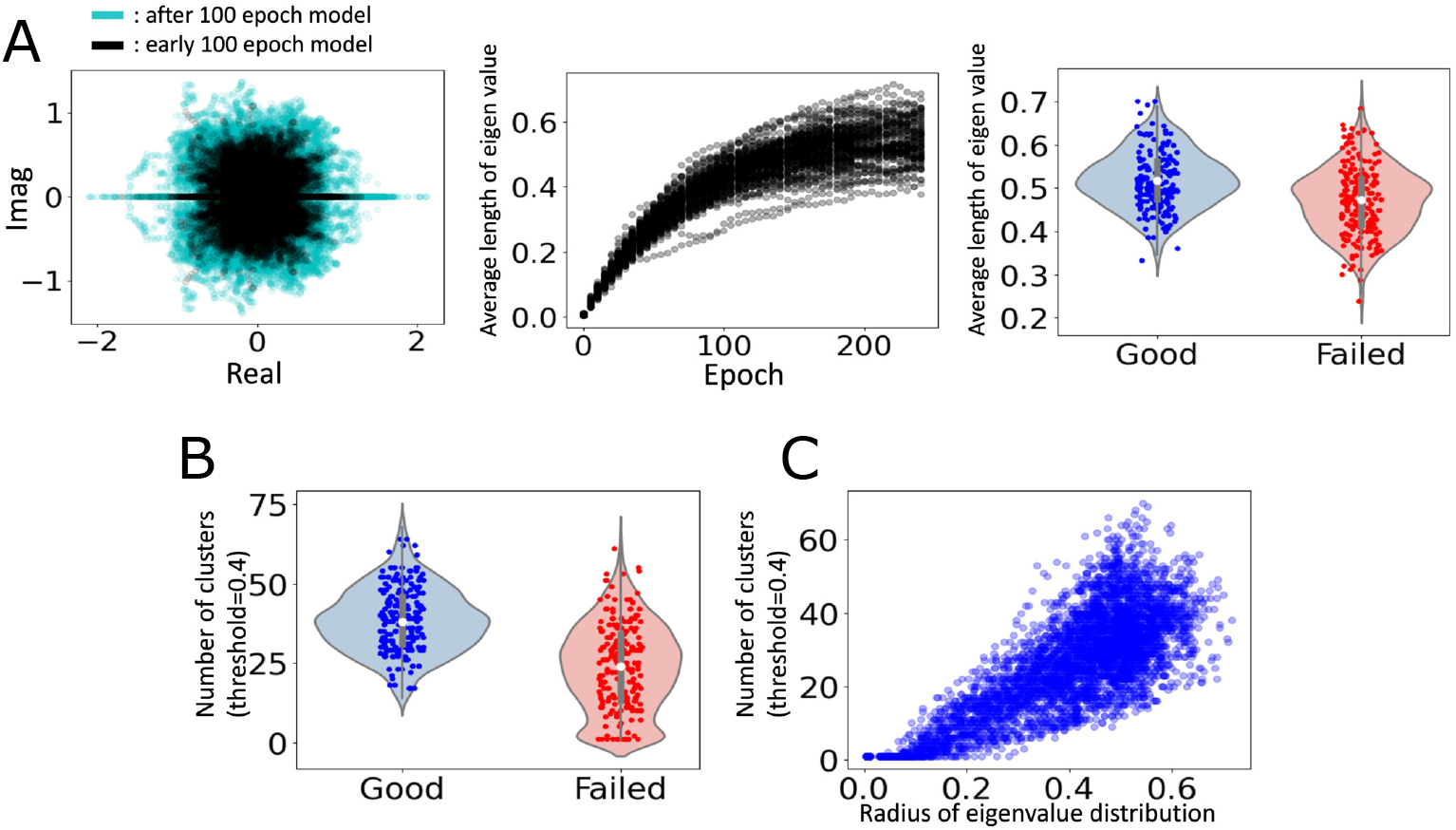
Diverse dynamical modes of successful models. **A**. Left: The eigenvalue distributions of recurrent connection weights of the PFC module are compared between early (black, epoch<100) and late (cyan, epoch >100) phases of training. Center: The evolution of the average radius of the eigenvalue distribution is plotted during learning. Right: Comparison of the mean eigenvalue radii between successful (blue) and failed (red) models (n=100 per group, *p*<0.01, ANOVA). **B**. The number of clustered activity patterns in the PFC module (hierarchical clustering, with threshold 0.4) is compared between successful and failed models. The analyzed period includes the movement phase (40 steps) and the delay period (80 steps). **C**. Correlations between the eigenvalue radius and cluster count are plotted for all trained networks.

We speculate that a broader eigenvalue distribution in the weight space confers greater flexibility in the choice of dynamical features available for learning representations. To assess this speculation, we quantitatively examined the repertoire of activity patterns in the Good and Failed models at each time point. The repertoire was quantified by temporally segmenting the activity of the PFC module in each model category and assessing the number of clusters identified through hierarchical clustering (Fig 6B). A model may be regarded as having a richer repertoire if more clusters emerge, giving a larger total cluster count. Successful models yielded more clusters than failed models for a given threshold for cluster detection. Furthermore, we observed a significant positive correlation between the cluster counts and eigenvalue spectra for the PFC modules of different model realizations (*r* =0.9510, *p<*0.001), implying that as eigenvalue distributions broaden, the granularity of information representation at each time point increases (Fig 6C). Altogether, these results confirm our speculation.

### Comparison with impaired 3-module models

We further explored the crucial role of the Re module in learning the present DNMS tasks by testing three other models with impaired or simpler network architectures. In the three-module model, the HPC module receives spatial information through an external input, and this crucial information is directly routed to the PFC module through HPC-to-PFC connections. However, the model also has an indirect pathway from HPC to PFC mediated by the Re module. We explored the roles of this pathway by eliminating either HPC-to-Re connections (UniHPC model) or Re-to-PFC connections (UniPFC model) (Fig 7A). Other connectivity structures and parameter values were unchanged. The number of trainable parameters in each model is listed in S1 Table.

**Fig 7.**
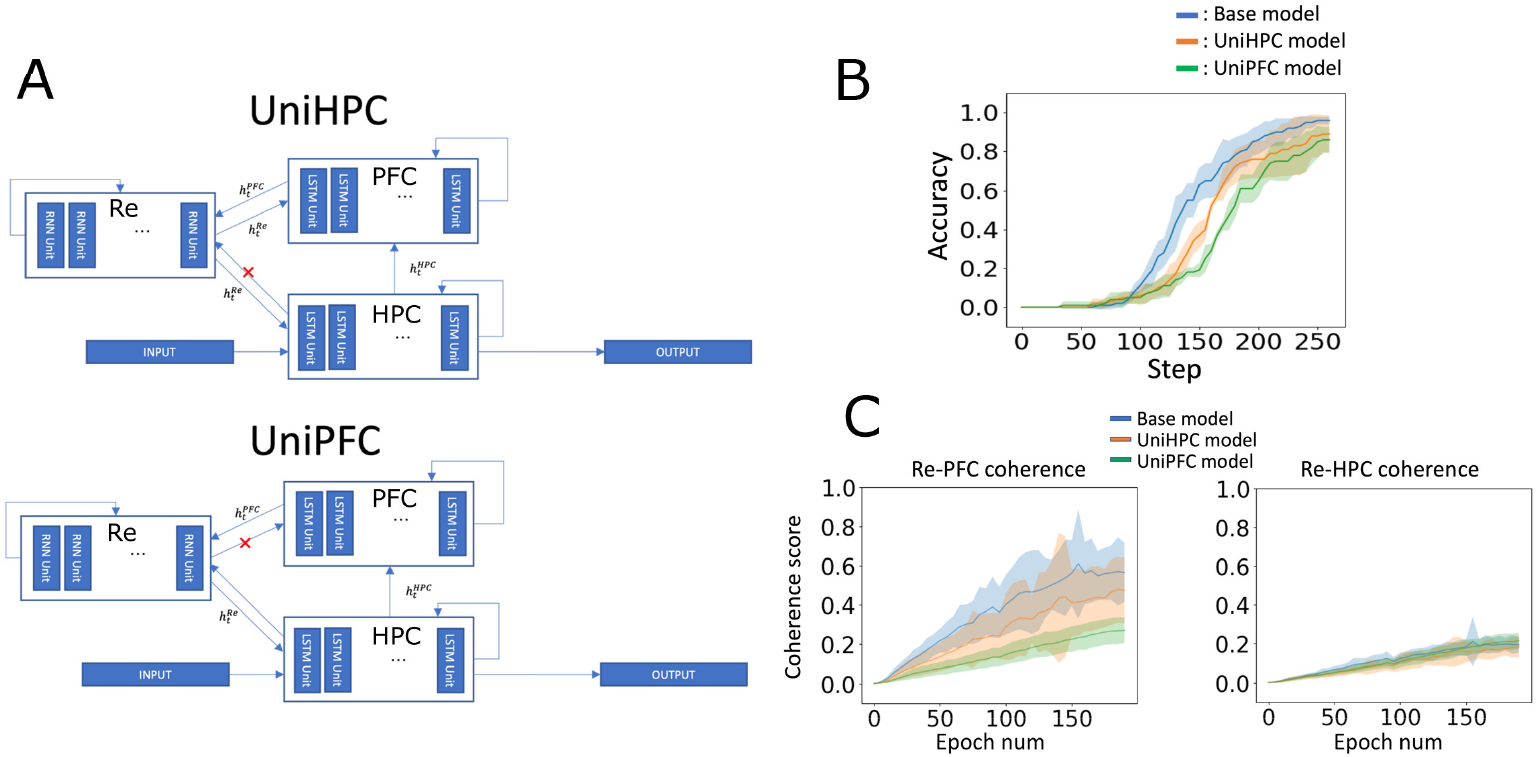
Performance of impaired models in learning. **A**. Two impaired three-module models were analyzed. The UniHPC model and UniPFC model lack HPC-to-Re projections (top) and Re-to-PFC projections (bottom), respectively. **B**. Learning performance in the H-maze DNMS task is shown for the original three-module model (blue), UniPFC model (red), and UniHPC model (green). **C**. The evolution of the average coherence scores (solid lines) during learning is shown for Re-PFC (left) and Re-HPC (right) module pairs in the three models. In B and C, shaded areas show SD.

Compared with the original model, the accuracy of the learned trajectories was significantly degraded in both impaired models, with the worst performance shown by the UniPFC model (Fig 7B). We then investigated how synchronization dynamics differed between the three models. The coherence between the HPC and Re modules did not exhibit large differences among the three models (Fig 7C, right). In contrast, the coherence score between the PFC and Re modules displayed marked differences and degraded according to the performance qualities of the three models, with the lowest score in the UniPFC model (Fig 7C, left). These results may be understood through the functional relationships of the three modules in these models. Although the UniHPC model lacks direct input from the HPC module to the Re module, it preserves connections from the RE module to both the HPC and PFC modules. The preserved connections enable the Re module to entrain the HPC and PFC modules, preventing a significant drop in the model’s performance. However, the UniPFC model lacks Re-to-PFC connections, presumably disenabling the entrainment of the PFC module by the Re module and degrading performance compared to the other models. Altogether, these results further demonstrate the crucial contribution of the Re module to navigating complex spatial environments in complex temporal contexts (e.g., working memory demands).

## Discussion

We investigated whether and how a neural network with three modules, representing the HPC, PFC, and a thalamic component (possibly Re), learns a complex memory-guided, spatial decision-making task, while paying attention to their divisions of labor. By analyzing activity patterns in each module, which was described by an LSTM unit with or without gating, we revealed that the HPC module forms attractor states that represent spatial information about a complex maze. The PFC module not only represents spatial information in a way that reflects the global task structure, but also acquires delay-period activity that captures the temporal and contextual information about the task. Our model suggests that the Re module is crucial for task execution by switching between different learned movements.

The dynamical features of the individual processing modules in our model align with various biological findings from the PFC-thalamic-HPC network. In vivo studies have shown that the reuniens enhances neural activity in the hippocampus [14, 29] and prefrontal cortex [22]. In spatial navigation, silencing the nucleus reuniens substantially reduced trajectory-dependent firing of CA1 neurons in the rat hippocampus [30]. It was recently shown that the lesions of the ventral midline thalamic nuclei impair the spatial representations encoded by CA1 place cells [31]. These results suggest that the activation of the ventral midline thalamic nuclei stabilizes hippocampal spatial representations. In our model, the Re module makes a significant contribution to the learning of stable task-related behavior, which is consistent with these experimental findings.

Both in experiments and our model, the ventral midline thalamic nuclei likely contribute to spatial decision-making by synchronizing PFC and HPC. The ventral midline thalamus increases performance measures in spatial working memory tasks that depend on interactions between the HPC and PFC [22]. Theta-rhythmic synchronization between PFC and hippocampus is crucial for success in alternating spatial navigation tasks [32] and is increased by interactions between the prefrontal cortex and ventral midline thalamus [33]. Other learning tasks, such as associative learning of eyeblink conditioning [34] and suppression of recalling extinguished fear memory [35], also rely on interactions between PFC, HPC, and the midline thalamic nuclei. As demonstrated in our model, owing to the bidirectional connections between the thalamus and the PFC/HPC, the thalamic nuclei occupy a unique position for effectively coordinating the activities of PFC and HPC. Indeed, the role of the Re module predicted in this model may also provide a mechanistic account for similar roles of other thalamic nuclei (e.g., the mediodorsal thalamic nucleus [36–38]) in memory-guided behaviors that recruit PFC and HPC.

Previous studies have employed spiking neural networks to simulate the structures and dynamics of brain regions, including thalamocortical interactions [39–41]. However, our model is based on rate-coding machine-learning units, which may sophisticate the ability of sequence learning of our model but limit its ability to replicate biological reality. An immediate consequence of this is that our model cannot tell whether the synchronization between the PFC, HPC, and Re occurs in the theta-frequency band. Indeed, synchronous neural activities have been observed in various oscillatory patterns, including theta [35], beta [24], gamma and theta-nested gamma oscillations [22], and sharp-wave ripples [42], in the PFC-thalamo-HPC network. A biologically more realistic model is necessary to address the frequency ranges of the inter-areal synchronization for spatial working memory tasks.

We implemented the Re module as an Elman-type recurrent network without gating. It learned fluctuating behavior with dual components essential for learning performance without hard-wired mechanisms to impose oscillations and multiple timescale dynamics. In our preliminary study, we found that the characteristic fluctuations largely disappeared if the Re module consisted of LSTM units, as in the other modules. This suggests that the fluctuating Re dynamics are not mere noise but reflect learned recurrent network dynamics, which LSTM likely suppresses with strong gating. Our results suggest that fluctuations in the Re dynamics maintain task-relevant representations that contribute to flexible state transitions in the entire triangular network.

Our model yields insights into the computational role and underlying mechanisms of the nucleus reuniens Re. This nucleus may serve as a site for integrating the task-relevant information stored in HPC and PFC via synchronization in different frequency bands. Synchronization can play multiple functional roles, including facilitation of effi-cient information transfer within and between local neural circuits. In our model, the slow and fast components contribute mainly to coordinating behavioral responses in accordance with the temporal structure of the task (e.g., the timing of movement initiation) or switching behavioral responses (e.g., right vs. left turns) according to decision rules, respectively. Similarly, frequency-specific synchronization may determine the context-dependent utilization of task-relevant information in the PFC-thalamo-HPC circuit under similar task structures. We speculate that our results are particularly relevant to theta–gamma coupling observed in Re activity. It has been suggested that Re regulates inter-areal communications between the hippocampus and prefrontal cortex by temporally coordinating activities in these regions [22, 24, 35, 42]. Whether and how the PFC-thalamo-HPC network establishes frequency-specific inter-regional connectivity to support different functional roles is open for further experimental and computational studies.

In summary, we constructed a triangular modular network model mimicking the prefrontal-thalamo-hippocampal circuit and explored how this network learns complex memory-guided spatial decision-making tasks that are also difficult for animals to learn. Our results suggest that the triangular network architecture formed by PFC, HPC, and thalamus can enhance the robustness of learning, especially for complex behavioral tasks.

## Supporting information

**S1 Fig.**
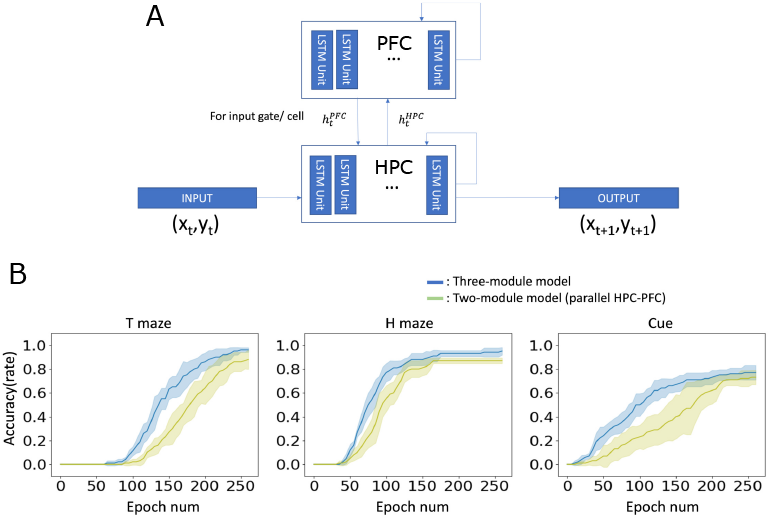
Comparison with a two-module model lacking the Re module. **A**. A modular network model without the Re module was trained on the T-maze, H-maze, and cue-guided DNMS tasks. The model had reciprocally connected HPC and PFC modules. **B**. Learning curves of the 3-module (blue) and 2-module (light green) models are plotted for the three DNMS tasks. All models achieved asymptotic performance exceeding 85% accuracy in the T-maze and H-maze tasks. Intriguingly, the three-module model outperformed the other models in learning speed in the H-maze and cue-based tasks, suggesting the advantage of the biologically inspired three-module architecture for navigating complex spatial environments in complex temporal contexts (e.g., working memory demands).

### S1 Table. Architectural parameters across network model variants

Parameter counts are presented for five neural network architectures: the proposed three-module model (Re), two-layer sequential LSTM model (Compare), two-module PFC-HPC model, and two unidirectionally connected variants (UniHPC and UniPFC). External weights represent connections from input layers to network modules; internal weights represent recurrent and inter-module connections; output weights connect to prediction layers.

## Methods and Materials

### Neural network model

In this study, all models were implemented in Python using PyTorch. We prepared two models: a three-module model and a two-module model. The three-module model consists of three modules: the HPC, PFC, and Re modules. The HPC and PFC modules are composed of LSTMs and have mutual connections internally. On the other hand, the Re module is constructed as an Elman network. The HPC module is connected to both the PFC and Re modules, the PFC module is connected to the Re module, and the Re module is connected to both the PFC and Re modules. Each module is composed of 20 units. The Re module influences the gating functions of the PFC and HPC modules, and inputs are provided to the Input gates and Cells of each module. External input is given to the HPC module, which generates output according to the task. The two-module model contains the HPC and PFC modules. Similar to the three-module model, it is composed of LSTMs and has mutual connections internally. The HPC module is connected to the PFC module, and inputs are provided to the Input gates and Cells of the PFC module. Each module is composed of 30 units. External input is given to the HPC module, which generates output according to the task (details provided in the task and training method descriptions below). This unit size was chosen to balance model complexity and training stability. While we also tested models with larger hidden sizes (e.g., 100 units), we observed no significant improvement in performance. In fact, training became more sensitive to hyperparameter settings (e.g., learning rate), and the overall learning stability decreased. Based on these observations, 20 units were adopted as a stable and efficient configuration for the current tasks.

**S1 Table.** Architectural parameters across network model variants.

| Model name | External weights | Internal weights | Output weights | Total parameters |
| --- | --- | --- | --- | --- |
| 3-module model | 4160 | 3600 | 40 | 7800 |
| Two module model (PFC-HPC) | 5640 | 7200 | 60 | 12900 |
| UniHPC model | 3760 | 3600 | 40 | 7400 |
| UniPFC model | 3360 | 3600 | 40 | 7000 |

The Re module was implemented as a simple Elman-type recurrent network without gating mechanisms, in contrast to the LSTM modules used for PFC and HPC. This architectural choice reflects the interpretation of the thalamus (Re) as a relay or modulatory hub that does not perform complex memory gating by itself. In preliminary tests, replacing the Re module with an LSTM significantly impaired learning efficiency without improving task performance. Although we do not include the comparison figures here due to space and time constraints, the Elman-type network yielded stable and consistent results across trials, and thus was adopted as a suitable architecture for this module. The Re module was connected only to the input gate and memory cell of each LSTM unit, reflecting a biological assumption that thalamic signals modulate input but are not themselves gated. Initially, we considered connecting Re only to the input gate, but this led to reduced task accuracy. Adding connections to the memory cell restored performance while preserving the intended role of Re. We deliberately avoided connections to the forget gate, both to minimize interference with temporal integration and because models using input+cell connections outperformed those using forget+cell configurations.

As comparative models, we also prepared the UniHPC model and UniPFC model. The UniHPC model is a version of the three-module model where the connection from the HPC module to the Re module is removed. All other settings are the same as in the three-module model. The UniPFC model is a version of the three-module model where the connection from the Re module to the PFC module is removed. All other settings are the same as in the three-module model.

An LSTM is a recurrent neural network model that first appeared in 1997 [25], and then many extend is developed. In [26], an LSTM unit was proposed, which comprises four gating functions. Each LSTM unit contains an input, output, and forget gate, as well as a cell. Input and output gates control the value of input or output. On the other hand, forget gate controls how the previous internal value of the unit affects the value of the next time step.

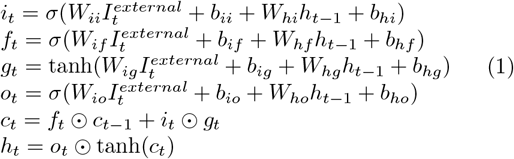

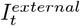, *h*_*t*_ are the input and hidden vector of the module. *c*_*t*_ is the internal vector. *i*_*t*_, *f*_*t*_, *g*_*t*_, *o*_*t*_ are the value of the input, forget, cell, and output gate at time step t.*W*_*ii*_ represents the weight matrix related to the input gate and input vector, *W*_*hi*_ is related to the input gate and hidden vector.*b* is the bias of each gate.*σ* and tanh is the activation function (sigmoidal and hyperbolic tangent).

For the Re module, we implemented an Elman network as follows:

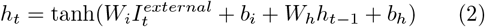

where 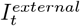 and *h*_*t*_ are the input and hidden state vectors at time step *t*, respectively. *W*_*i*_ the input-to-hidden weight matrix, *W*_*h*_ is the hidden-to-hidden (recurrent) weight matrix, and *b*_*i*_ and *b*_*h*_ are the corresponding bias vectors. tanh denotes the activation function (hyperbolic tangent).

### Context-dependent task

The delayed non-match task is a spatial navigation task performed in a T-maze or H-maze-like environment. In this task, the subject, such as a rat, is required to reach one side of the maze and then return to the initial position. After a delay, the subject is required to go to the other side of the maze to receive a reward. This task requires working memory to remember which arm was visited and to understand the rules of the environment. The T-maze is represented in X-Y coordinates. The X values range from (0.25–0.75) and the Y values from (0.0–0.5), with the initial position set at (0.5, 0). Two types of trajectories are prepared: one route leading to the right arm and another to the left arm. The goal points are set at (0.75, 0.5) for the right arm and (0.25, 0.5) for the left arm. Similarly, the H-maze is also represented in X-Y coordinates. It contains four arms: lower right (A), upper right (B), upper left (C), and lower left (D). Four types of trajectories are prepared, corresponding to visiting two arms following the rules A→D, B→C, C→A, and D→B. The goal points are set at (0.75, 0.25) for A, (0.75, 0.75) for B, (0.25, 0.25) for C, and (0.25, 0.75) for D.

For quantitative assessment of functional specialization between modules, we employed task-specific analytical approaches optimized for different purposes. T-maze data was used for correlation methodology illustration due to its simple left-right trajectory structure that clearly demonstrates the relationship between spatial differences and neural representations. H-maze data was used for temporal analysis because its four distinct trajectory patterns (A→D, B→C, C→A, and D→B) provide more comprehensive statistical power for examining how spatial and contextual information segregation develops across extended training periods.

Experiments were also conducted on tasks where a cue was input. The basic trajectory remains the same, but in this case, cue information is added as external input. The timing of the added cues occurs at the very beginning of the trajectory and at the end of the delay period, right before the subject starts moving after the delay. Therefore, in this task, the subject does not need to learn the length of the delay but rather needs to learn the behavior corresponding to the cue information. A value of 1 is input when the cue is provided, and a value of 0 is input at all other times.

The trajectory consists of Movement periods 1 and 2, and a Delay period. Movement periods 1 and 2 are the times when the arms are visited, and the Delay Period is when the subject waits at the initial position. The trajectory length in the T-maze is 100 steps in total: 60 steps for the Delay Period and 20 steps each for Movement periods 1 and 2. In the H-maze, the total trajectory length is 160 steps: 80 steps for the Delay Period and 40 steps each for Movement periods 1 and 2. In the Cue task for the T-maze, the lengths of Movement periods 1 and 2 remain at 20 steps each, but the length of the Delay Period is randomly set between 40 and 80 steps.

Each model was trained for 8,000 epochs with a batch size of 20. Each epoch corresponded to one training episode, which included a single target trajectory.

For the T-maze and cue task, successful performance was defined as alternating between two maze arms, determined by crossing x-coordinate thresholds (Figure 1B : x < 0.2 on the left arm; x > 0.7 on the right arm) at least three times in a correct sequence. For the H-maze task, the model needed to reach specific arm coordinates in the prescribed order of A→D, B→C, C→A, and D→B (Figure 1C). Arm visiting was defined by the trajectory crossing threshold coordinates (x = 0.2 or x = 0.7 for horizontal boundaries, and y = 0.3 or y = 0.6 for vertical boundaries) and reaching within a Euclidean distance of 0.15 from the corresponding goal points at (0.75, 0.25), (0.75, 0.75), (0.25, 0.25), and (0.25, 0.75) for arms A, B, C, and D respectively. This distance threshold was set based on the arm length of 0.25. A model was classified as “Good” (well-trained) when it successfully completed the required alternating sequence for the T-maze or the prescribed arm-visiting order for the H-maze. A model was classified as “Failed” if it became stuck in the first-visited arm or showed repetitive, non-directional movement patterns. Accuracy was calculated as the percentage of successful models across 100 independent training runs.

### Training method

The learning method was conducted using Back Propagation Through Time (BPTT) [43]. For each training episode, 50 time points were randomly sampled from the full target trajectory. For each sampled time point *t*, the model was trained to reproduce the trajectory from the initial state up to time *t*. The mean squared error (MSE) between the model’s predicted position at time *t* and the corresponding target was used as the loss function. This process was repeated for all time points in order of increasing *t*, resulting in 50 weight updates per episode via BPTT. Optimization of the weight matrix was performed using BPTT with ADAM optimizer [44] with a learning rate of 0.0002 and weight decay of 1e-8. We initialized network weights with random values sampled from a Gaussian distribution N(0, 0.01). Hidden states were also randomly initialized from N(0, 0.01) at the start of each trajectory. The input to the network consisted of position coordinates (x, y) for the T-maze and H-maze tasks, and position coordinates plus a binary cue signal (x, y, cue) for the cue-applied task, updated at every time step. To reiterate, one weight update is performed for each of the 50 sampled time points, resulting in 50 updates per episode. To ensure statistical reliability, we trained 100 independent models for each architecture configuration.

### Statistical analysis

Principal component analysis (PCA) was employed as an exploratory dimensionality reduction method to visualize and compare population-level activity across modules and task phases. The goal of this analysis was not to identify a minimal generative basis, but to extract low-dimensional structure that captures dominant patterns of activity relevant to task performance. When comparing the differences between two trajectories, the outputs obtained from these trajectories were combined into a single time-series data set. PCA was then applied to reduce the dimensionality, after which the data was divided back into its original length for further analysis. To determine the number of components to retain, principal components were included until they collectively explained approximately 60-70% of the total variance in neural activity. This criterion provided a balance between capturing dominant task-related structure and avoiding higher-order components that primarily reflected noise or fluctuations. Statistical significance was assessed using ANOVA.

### Coherence score

Each module exhibits periodic activity. To investigate the coherence between the Re module and the PFC/HPC modules in terms of this activity, a short-time Fourier transform with a window width of 60 steps was applied to each module, and the resulting spectrograms were analyzed. The cross-spectrum between the Re-HPC and Re-PFC modules was calculated based on the obtained spectrograms. The coherence score was determined by calculating the difference between the maximum and minimum values of the time series data of the averaged similarity across frequencies. This allowed us to assess the similarity of frequencies corresponding to periodic activity between the modules. The coherence score was calculated for the model at each epoch to observe how it changed with learning.

### Eigenvalue

To explore whether characteristics of the model could be extracted from the weight matrix, the eigenvalues of the weight matrix were examined. The matrices of interest were the weight matrices of the input gate and cell in the internal connections of the PFC/HPC, which were multiplied together. The resulting matrix size was (20,20) * (20,20) = (20,20). The eigenvalues obtained from this matrix were plotted in a scatter plot to observe their distribution in the eigenvalue space. The distribution’s radius was plotted for each epoch, with the radius being the average distance of each eigenvalue from the origin.

### Clustering

To confirm this assumption, I examined the similarity of activity patterns at each time point for each model. Activity similarity at each time step was quantified using dendrogram-based hierarchical clustering, which groups activity patterns according to their pairwise similarity structure. Pairwise distances between PFC activity patterns were computed using Euclidean distance, and clusters were iteratively merged using Ward’s linkage method.

This procedure yields a dendrogram that summarizes hierarchical relationships among activity patterns without requiring an a priori specification of the number of clusters. A distance threshold was applied to the dendrogram to estimate the approximate number of distinguishable clusters under this similarity metric. This thresholding was not intended to define discrete or categorical neural states, but rather to provide a coarse characterization of representational separation in the PFC module. A threshold of *θ* = 0.4 was used for examining PFC clusters, and *θ* = 0.2 for HPC clusters. The general characteristics remained consistent even when the value of *θ* was altered. If similarities are high, fewer clusters should emerge, reducing the total cluster count. The analyzed activities include the movement period(40 steps) and the delay period(80 steps).

## Data Availability

All training data used in this study were synthetically generated using the simulation python scripts provided in the GitHub (https://github.com/oist-ncbc/takaku_fukai_2026) repository. Because the datasets are generated stochastically, the exact instances cannot be perfectly reproduced; however, statistically equivalent data can be regenerated by running the provided scripts with the same configurations. The trained model checkpoints are available via Zenodo. The repository contains all source code necessary to retrain the network models and reproduce the reported analyses.

## Acknowledgments

Thanks for the lab member, Prof. Gordon W. Arbuthnott, Prof. Kazumasa Tanaka, and Prof. Tomoki Fukai as kind supervision.

## Declaration of generative AI and AI-assisted technologies in the manuscript preparation process

During the preparation of this work the authors used Grammarly and Claude in order to assist with the organization and language editing of the manuscript. After using this tool/service, the authors reviewed and edited the content as needed and take full responsibility for the content of the published article.

